# Cell Painting transfer increases screening hit rate

**DOI:** 10.1101/2022.10.24.513564

**Authors:** Ethan Cohen, Maxime Corbe, Cláudio A. Franco, Francisca F. Vasconcelos, Franck Perez, Elaine Del Nery, Guillaume Bollot, Auguste Genovesio

**Author notes:** correspondence should be addressed to AG.

## Abstract

Drug discovery uses high throughput screening to identify compounds that interact with a molecular target or that alter a phenotype favorably. The cautious selection of molecules used for such a screening is instrumental, and is tightly related to the hit rate. In this work, we wondered if Cell Painting, a general purpose image-based assay, could be used as an efficient proxy for compound selection, thus increasing the success rate of a specific assay. To this end, we considered Cell Painting images with 30,000 molecules treatments, and selected compounds that produced a visual effect close to the positive control of an assay, by using the Frechet Inception Distance. We then compared the hit rates of such a preselection with what was actually obtained in real screening campaigns. As a result, Cell Painting would have permitted a significant increase in the success rate and, even for one of the assays, would have allowed to reach 80% of the hits with ten times fewer compounds to test. We conclude that images of a Cell Painting assay can be directly used for compound selection prior to screening, and we provide a simple quantitative approach in order to do so.

## Introduction

Target-based high throughput screening is currently the main approach to drug discovery. It consists of identifying active chemical compounds that interact with a pre-identified target through automated parallelized experimental tests (Macarron *et al*., 2011; Swinney and Anthony, 2011). Another approach that has gained in popularity, especially in seeking first-in class therapeutic drugs, is phenotypic screening, during which one seeks compounds that modulate a phenotype of interest (Kotz, 2012; Zheng, Thorne and McKew, 2013). To this end, a library of compounds is selected in the hope that a part of these, defined as “hits”, will reproduce or approach the phenotype obtained with a positive control tool treatment.

One of the main issues that concerns all types of molecular screens is the selection of the compound library to be tested, as this will have an impact on the final hit rate. Given the size of the chemical space, which is in the order of 10^60^, a random selection of compounds is largely suboptimal. Various methods have been proposed to reduce the attrition rate of a screening, through a better preselection of compounds to test (Waring *et al*., 2015). Furthermore, specific libraries have been designed, such as the Dundee Kinase Inhibitor library to address specific modulation types (Brenk *et al*., 2008).

Cell Painting (CP) is a phenotypic image-based assay developed at the Broad Institute that offers to capture high-level and general information from images of cells under perturbations, through labeling of important organelles (Bray *et al*., 2016). These organelles range from the nuclei, to nucleoli, mitochondria, actin and tubulin networks. On top of CP’s ability to monitor the state of the cells, the images of a large U2OS cell painting screen containing 30K compounds were made publicly available (Bray *et al*., 2017). The community also expects the release of CP images of 120,000 compound treatment by the JUMP consortium in late 2022 (Chandrasekaran *et al*., 2022).

In this work, we wondered whether CP coupled with transfer learning could provide a good proxy to efficiently pre-select active compounds for a specific assay. To this end, we place ourselves in a real case scenario where we evaluate the hit rate we would have obtained on high-content screening campaigns we performed, had we used cell painting prior to screening to select 5%, 10% or 15% of the compounds.

## Results

### Transfering hits from cell painting through Inception

To evaluate the preselection gain that cell painting could offer in practice, we scanned recently performed in-house Compound Screening campaigns (CPDS at Institut Curie, Biophenics). For each assay, we considered all compounds that were both in the considered screen and were part of the 30k compound tested in the CP assay introduced in Bray et al (Bray *et al*., 2017). We considered only the assays that had a positive control and at least 380 compounds and 5 hits in common, which ended up in the selection of 3 screens. For each of these 3 screens, we then ranked the compounds by decreasing similarity with our positive control using the CP assay. To measure similarity between images of cells perturbed by two compound treatments in the CP assay, we used the Frechet Inception Distance (FID, see Methods) (Heusel, Ramsauer and Unterthiner, 2017). In short, computing the FID between two conditions consists in vectorizing all the images from each condition through a pretrained convolutional network (Inception) and computing the Frechet distance between these high-dimensional sample feature distributions (Szegedy *et al*., 2015). We then examined how many of the hits we actually obtained during the screening campaign would have been ranked in the first 5%, the first 10% and the first 15% of this list. Furthermore, to assess the significance of these results, we performed a Fisher exact test (Fisher, 1922). **Table 1** reports results we obtained for each screen.

**Table 1:** Hit prediction using Cell Painting on 3 recent screening campaigns - The first column lists the compound screening campaign (CPDS). The second column indicates the number of compounds tested in the screen that were also found in the Cell Painting assay. The third column indicates, among those compounds, how many were selected as hits in the considered screen. The three remaining columns indicate the ratio of hits obtained in the Top 5, 10 and 15% of the ranked list of compounds. The p-value of an exact Fisher test is reported in parentheses.

| Screens | Compounds also in CP | Hits also in CP | Top 5% | Top 10% | Top 15% |
| --- | --- | --- | --- | --- | --- |
| CPDS#1 | 400 | 5 | 2/5 (2.1e-2) | 4/5 (4. e-4) | 4/5 (2.1e-4) |
| CPDS#2 | 380 | 32 | 14/32 (3.7e-13) | 19/32 (1.2e-13) | 19/32 (1.0e-9) |
| CPDS#3 | 360 | 38 | 7/38 (1.1e-3) | 10/38 (1.8e-3) | 15/38 (7.5e-5) |

### Robustness to variation of experimental settings

In any primary screen, a dedicated protocol is followed. It typically comprises many parameters such as cell line, cell seeding, incubation time, concentration or labeling. This protocol varies from screen to screen depending on the end goal and the Cell Painting assay presented in Bray et al. doesn’t escape this rule. The consequence is that the same perturbation in two different assays does not necessarily produce the same phenotypes. In fact we observed that cell phenotypes can be significantly different even with a similar compound treatment (**Figure 1A** and **B**). However, interestingly, these variations in protocols didn’t prevent the prediction from being highly significant. The reason may be that while a difference in phenotype is obvious for the same compound across screens, it doesn’t prevent the control and the hits to look similar in each individual screen. For instance, **Figure 1A** and **C** display the Cell Painting images of the positive control condition Brefeldin A for the CPDS#2, and Piperlongumine, a compound correctly predicted as a hit using FID. The images of phenotypes in Cell Painting expectedly look similar as Piperlongumine was selected by this means. However, while these phenotypes in CP both look dissimilar with the phenotypes in the CPDS#2 (probably due to the difference in incubation time), they seem to again match when considering CPDS#2 only.

**Figure 1:**
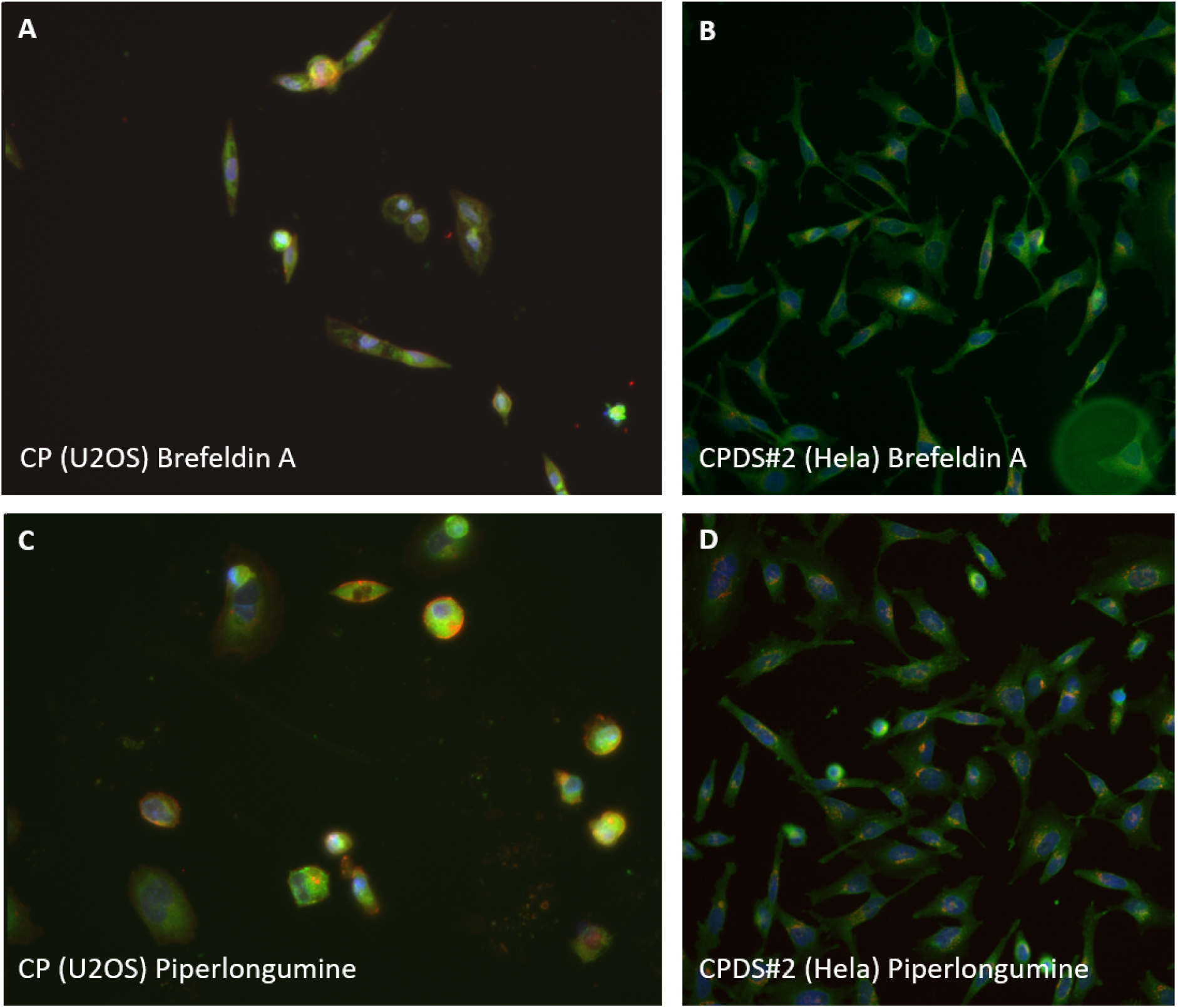
**A -** effect of Brefeldin A treatment on the Cell Painting assay (5μM, U2OS, incubation time: 24h) **B**- effect of Brefeldin A treatment on the CPDS#2 (10μM, Hela, incubation time: 120 min) **C**- effect of Piperlongumine on the Cell Painting assay, a compound selected as close to Brefeldin A using FID and independently as a hit in CPDS#2. **D**- effect of Piperlongumine on the CPDS#2. A and B produce dissimilar phenotypes, but A is close to C, and B is close to D.

### Robustness to the positive control relevance

CPDS#1 and 2 were designed to monitor the trafficking of fluorescently labeled reporter proteins out of the ER using the Retention Using Selective Hooks (RUSH) assay (Boncompain and Perez, 2013). In such screens we seek for compounds that reproduce a specific phenotype, precisely defined by the positive control compound Brefeldin A (BFA). On the other hand, CPDS#3 is a cell survival screen where the goal is to identify toxic compounds, but no specific positive control was used to this end. Instead, hits were formerly selected based on cell count. We then applied the following strategy: we arbitrarily chose two cytotoxic compounds (Lovastatin and Fluvastatin) as positive controls in Cell Painting assay in order to select a list of possibly toxic compounds (Buchou *et al*., 2022). Interestingly, the results for CPDS#3 displayed in **Table 1** remain highly significant, which suggests that - at least for such an obvious phenotype such as cell death - it seems sufficient to grab a selection of candidates that loosely reproduces the effect of a toxic compound, to perform better than a random selection of compounds.

## Discussion

In this work, we wondered if CP could be used in practice to select an efficient list of compounds to be tested before a screening campaign. To this end, we propose to simply use the Frechet Inception Distance as a metric in a large Cell Painting assay, to select compounds likely to reproduce the phenotype obtained with a positive control in a specific assay. Using already achieved screening campaigns, we quantitatively demonstrated that this strategy would have allowed, in practice, to drastically reduce the number of molecules to screen, while consistently improving hit rate. Concretely, screening only 10% of compounds using this strategy would allow us to pick up from 25% to up to 80% of the hits, depending on the assay. Overall, it seems to be constantly beneficial to use Cell Painting for compound selection prior to screening, rather than using a compound library directly.

Furthermore, our results suggest that, while phenotypes produced by a given compound could largely vary from CP to a specific screen, due to the variation in experimental settings, e.g. cell model, labeling, compound concentration and incubation time, phenotype similarities of two compound treatments in cell painting was likely reproducible in a specific screen. Finally, our results suggest that, at least when the phenotype of interest is cell death, it is enough to arbitrarily select a few toxic compounds as positive controls, to obtain a compound selection that is more likely to lead to hits than random.

Importantly, the overlap between our compound library and the libraries used in Bray et al. reached about 400 compounds at best. In consequence, some highly ranked compounds in the Cell Painting assay could not be observed in practice in our specific screens. Therefore we anticipate that the hit ratio obtained using the suggested approach could be significantly higher in a real case scenario, when used for future screens where all compounds close to a positive control in Cell Painting could be tested. We also anticipate that this approach, straightforward to use, will also largely benefit from the soon to be released CP-JUMP dataset by the Broad Institute that is expected to comprise more than 120,000 compound treatments.

## Methods

### Screening assays

We performed an exhaustive search in the assays that were previously screened in house. We computed the intersection of compounds used in each screen and the 30k compounds tested in the CP assay introduced in Bray et al (Bray *et al*., 2017). We selected those assays that comprise the positive control and more than 3 hits in this intersection. This filtering led to only 3 assays among 60, mostly because the positive control in our assays were not frequently part of the CP assay. In some other cases, it was due to the discrepancies between the compound libraries leading to a small overlap.

The 3 compound drug screening campaigns (CPDS) retrieved this way were performed using 1,600 compounds obtained from Prestwick (1,280 off-patent small molecules, mostly approved drugs from FDA, EMA, and other agencies and a set of 320 phytochemical compounds), all tested at 10 μM.

For CPDS#1, A549 cells stably expressing the GFP-ACE2 RUSH reporter (Boncompain and Perez, 2013) were seeded in 384-wp (Viewplate 384, Perkin Elmer) for 24h, treated with compounds for 90 min, then subsequently treated with 40 μM of biotin for 60 min.

For CPDS#2 screen, HeLa cells stably expressing the GFP-VEGF RUSH reporter were seeded in 384-wp for 24h, treated with compounds for 120 min, then subsequently treated with biotin as previously described.

In both previous screens, RUSH reporters are retained in the endoplasmic reticulum (ER) through a streptavidin-based interaction that is relieved by biotin addition to cells. Brefeldin A (BFA) is added to the screen as a positive control of ER retention.

For CPDS#3 screen, we employed the Ewing sarcoma A673 cells with compounds being incubated for 24h, as previously described (Buchou *et al*., 2022).

Image acquisition was performed after fixation of cells with 4% formaldehyde solution and nuclei staining with 0.2 μg/mL of DAPI using the INCell 2200 automated widefield system (GE Healthcare, USA) at a 20X magnification (Nikon 20X/0.45, Plan Apo, CFI/60).

### Image-based sample similarity

To measure similarity between images of cells perturbed by two compound treatments in the CP assay, we used the Frechet Inception Distance (FID)(Heusel *et al*., 2017). FID was primarily designed to compare distributions of synthetic images generated by a generative model with real images used to train the model. Briefly, it consists in passing all images through an Inception v3 network pre-trained on ImageNet, then computing the squared Wasserstein metric between the two distributions approximated as multivariate Gaussian(Szegedy *et al*., 2015).

### Rank and statistical test

After ranking all compounds by decreasing order of similarity with a positive control using FID, we examined the fraction of hits we would have obtained had we decided to screen only the first 5%, the first 10% or the first 15% of the most similar compounds in CP. After this, we tested the significance of each of these ratios compared to the total ratio of hits. To this end we performed an exact Fisher test that computes a p-value, using the hypergeometric law, which is the exact probability to obtain a ratio equal or more extreme than the observed ratio.

## Acknowledgments

We thank Olivier Delattre (U830/DEPICT, Institut Curie) for providing access to the screening data. This work was supported by ANR–10–LABX–54 MEMOLIFE and ANR–10 IDEX 0001–02 PSL* Université Paris, ANRT CIFRE and was granted access to the HPC resources of IDRIS under the allocation 2020-AD011011495 made by GENCI. FFV was supported by a postdoctoral researcher contract from FCT (CEECIND/04251/2017).

